# A new role for histone demethylases in the maintenance of plant genome integrity

**DOI:** 10.1101/2020.03.02.972752

**Authors:** Javier Antunez-Sanchez, Matthew Naish, Juan Sebastian Ramirez-Prado, Sho Ohno, Ying Huang, Alexander Dawson, Deborah Manza-Mianza, Federico Ariel, Cecile Raynaud, Anjar Wilbowo, Josquin Daron, Minako Ueda, David Latrasse, R. Keith Slotkin, Detlef Weigel, Moussa Benhamed, Jose Gutierrez-Marcos

**Affiliations:** School of Life Science, University of Warwick, Coventry CV4 7AL, UK; Institute of Plant Sciences Paris Saclay, Bureau 1.54 - Bâtiment 630 - Rue Noetzlin, 91405 Orsay Cedex, France; Department of Molecular Biology, Max Planck Institute for Developmental Biology, Tubingen, Germany; Department of Molecular Genetics, The Ohio State University, Columbus, OH, USA; Institute of Transformative Bio-Molecules, Nagoya University, Furo-cho, Chikusa-ku, Nagoya, Aichi 464-8601, Japan; Donald Danforth Plant Science Center, St. Louis, MO, USA; Graduate School of Agriculture, Kyoto University, Kitashirakawa Oiwake-cho, Sakyo-ku, Kyoto, 606-8502; Division of Biological Science, Graduate School of Science, Nagoya University, Furo-cho, Chikusa-ku, Nagoya, Aichi 464-8602; Division of Biological Sciences, University of Missouri, Columbia, MO, USA

**Keywords:** Arabidopsis, Chromatin, DNA methylation, Epimutation, Transposon

## Abstract

Histone modifications deposited by the Polycomb repressive complex 2 (PRC2) play a critical role in the control of growth, development and adaptation to environmental fluctuations in most multicellular eukaryotes. The catalytic activity of PRC2 is counteracted by Jumonji-type (JMJ) histone demethylases, which shapes the genomic distribution of H3K27me3. Here, we show that two JMJ histone demethylases in Arabidopsis, EARLY FLOWERING 6 (ELF6) and RELATIVE OF EARLY FLOWERING 6 (REF6), play distinct roles in H3K27me3 and H3K27me1 homeostasis. We show that failure to reset these chromatin marks during sexual reproduction results in the inheritance of epigenetic imprints, which cause a loss of DNA methylation at heterochromatic loci and transposon activation. Thus, Jumonji-type histone demethylases in plants contribute towards maintaining distinct transcriptional states during development and help safeguard genome integrity following sexual reproduction.

## Introduction

In eukaryotes, chromatin accessibility is modified by DNA methylation, the covalent modification of histone proteins and the deposition of histone variants. These epigenetic modifications allow the establishment of specific transcriptional states in response to environmental or developmental cues. While in most cases environmentally-induced chromatin changes are transient, epigenetic changes induced during development are often stably inherited through mitotic divisions, so that cell identity is maintained and individual cells or tissues do not revert to previous developmental states. A key chromatin modification implicated in these responses is the post-translational modification of histone tails, which are associated with active or inactive transcriptional states (Kouzarides, 2007). Among these, the methylation of lysine 9 of histone H3 (H3K9me2) and H3K27me1 has been associated with the repression of transposable elements (TEs) in constitutive heterochromatin, whereas methylation in others, including H3K27me3, has been associated with the repression of genes in euchromatic genome regions (Berger, 2007, Pfluger & Wagner, 2007). The latter is deposited by PRC2 and plays a crucial role in development in most multicellular eukaryotes (Laugesen, Hojfeldt et al., 2019). In plants, this modification is found in approximately one quarter of protein-coding genes and is dynamically regulated during growth and development (Lafos, Kroll et al., 2011, Roudier, Ahmed et al., 2011, Zhang, Germann et al., 2007). The activity of PRCs is counterbalanced by JMJ demethylases, which catalyze the specific removal of H3K27me3 (Liu, Lu et al., 2010). In Arabidopsis, five histone demethylases [RELATIVE OF EARLY FLOWERING 6 (REF6); EARLY FLOWERING 6 (ELF6); JUMONJI 13 (JMJ13); JUMONJI 30 (JMJ30); and JUMONJI 32 (JMJ32)] have been implicated in the demethylation of H3K27 (Crevillén, Yang et al., 2014, Gan, Xu et al., 2014, Lu, Cui et al., 2011). These proteins are thought to mediate the temporal and spatial de-repression of genes necessary for a wide range of plant processes such as flowering, hormone signaling, and the control of the circadian clock. Inactivation of *REF6* results in the ectopic accumulation of H3K27me3 at hundreds of loci, most of them involved in developmental patterning and environmental responses (Lu et al., 2011, Yan, Chen et al., 2018). It has been proposed that REF6 is recruited to a specific sequence motif thought its zinc-finger domains (Cui, Lu et al., 2016, Lu et al., 2011), however others have shown that it is also recruited by specific interactions with transcription factors (Yan et al., 2018). Moreover, it has been shown that the affinity of REF6 to chromatin is hindered by DNA methylation, which could explain why its activity is primarily found at euchromatic loci (Qiu et al 2019).

Previous studies proposed that REF6 acts redundantly with ELF6 and JMJ13 to restrict the accumulation of H3K27me3 in gene regulatory regions thereby unlocking tissue-specific expression (Yan et al., 2018). Importantly, REF6, ELF6, JMJ30 and JMJ32 appear to specifically remove methyl groups from H3K27me3 and H3K27me2 but not from H3K27me1 (Crevillén et al., 2014, Gan et al., 2014, Lu et al., 2011). Previous investigations have shown that H3K27me1 in Arabidopsis is associated with constitutive heterochromatin, where it is deposited by ARABIDOPSIS TRITHORAX-RELATED PROTEIN5 (ATXR5) and ATRX6 (Jacob, Feng et al., 2009, Jacob, Stroud et al., 2010). However, several studies in mammals and plants have shown that H3K27me1 is also found in euchromatin (Fuchs, Jovtchev et al., 2008, Jacob et al., 2009, Vakoc, Sachdeva et al., 2006). The presence of H3K27me3 in euchromatin is thought to be actively re-set during sexual reproduction – a view supported by studies in Arabidopsis showing that ELF6, REF6 and JMJ13 are necessary to reset and prevent the inheritance of this epigenetic mark by the offspring (Crevillén et al., 2014, Liu, Feng et al., 2019, Zheng, Hu et al., 2019). However, to what extent these epigenetic imprints are reset during sexual reproduction remains unknown.

Here, we show that the histone demethylases REF6 and ELF6 play distinct roles in the demethylation of histones in Arabidopsis, and that REF6 is a major player in the deposition of H3K27me1 in active chromatin. We found that failure to reset H3K27me3 marks during sexual reproduction results in the inheritance of these epigenetic imprints even in the presence of fully functional histone demethylases. The ectopic inheritance of H3K27me3 is associated with the loss of DNA methylation at heterochromatic loci leading to activation of TEs. Moreover, we found that genetic and epigenetic mutations arising in histone demethylase mutants are stably inherited over multiple generations and result in pleiotropic developmental defects. Collectively, our work has uncovered a hitherto unrecognized role for histone demethylases in maintaining the genetic and epigenetic stability of plants.

## Results

### Arabidopsis REF6 and ELF6 play distinct roles in H3K27me3 homeostasis

The deposition of H3K27me3 by PRCs correlates with transcriptional repression both in plants and animals. The dynamic regulation of this epigenetic mark enables the reactivation of genes primarily implicated in developmental programs; thus, any disruption to these regulatory networks results in major developmental aberrations (Kassis, Kennison et al., 2017, Lewis, 1978, Molitor, Latrasse et al., 2016). The demethylation of H3K27me3 has been linked to the enzymatic activity of five JMJ-type proteins, which act antagonistically to SET-domain histone methyltransferases from the PRC2 complex (Yan et al., 2018). In order to gain further knowledge about these processes we investigated the function of two sequence-related histone demethylases, ELF6 and REF6, in Arabidopsis. We isolated a loss-of-function insertion located in the sixth exon of *REF6* (*ref6-5*) and a targeted CRISPR/Cas9 deletion to the first exon of *ELF6* (*elf6-C*) (Supplemental Fig. S1). Similar to previous reports, we found that our *elf6-C* plants displayed an early flowering phenotype, characterized by a reduced number of rosette leaves at bolting. Conversely, *ref6-5* plants displayed a late flowering phenotype and an increased number of rosette leaves at bolting stage (Fig 1A and Supplemental Fig. S2A-B). Moreover, *elf6-C/ref6-5* double mutant plants displayed a dwarf phenotype, increased number of petals and pleiotropic defects in leaf morphology, such as serrations and downward curling (Fig 1A and Supplemental Fig. S2C-D). Notably, the phenotypes observed in *elf6-C/ref6-5* have not been previously reported for double mutants of these histone demethylases (Yu, Li et al., 2008), which could be explained by the fact that only partial-loss-of-function mutations had been used previously (Yan et al., 2018). Further phenotypic analysis revealed that *elf6-C/ref6-5* plants displayed a reduction in silique length (Fig 1B), thus suggesting that these mutations may affect plant fertility. Microscopy analysis of developing seeds revealed that while embryo development in *elf6-C* was normal, seeds from *ref6-5* and *elf6-C/ref6-5* contained embryos that displayed patterning defects (Fig 1B). However, these embryonic abnormalities did not give rise to any noticeable changes in seed germination rates (Supplemental Fig. S2E).

**Figure 1.**
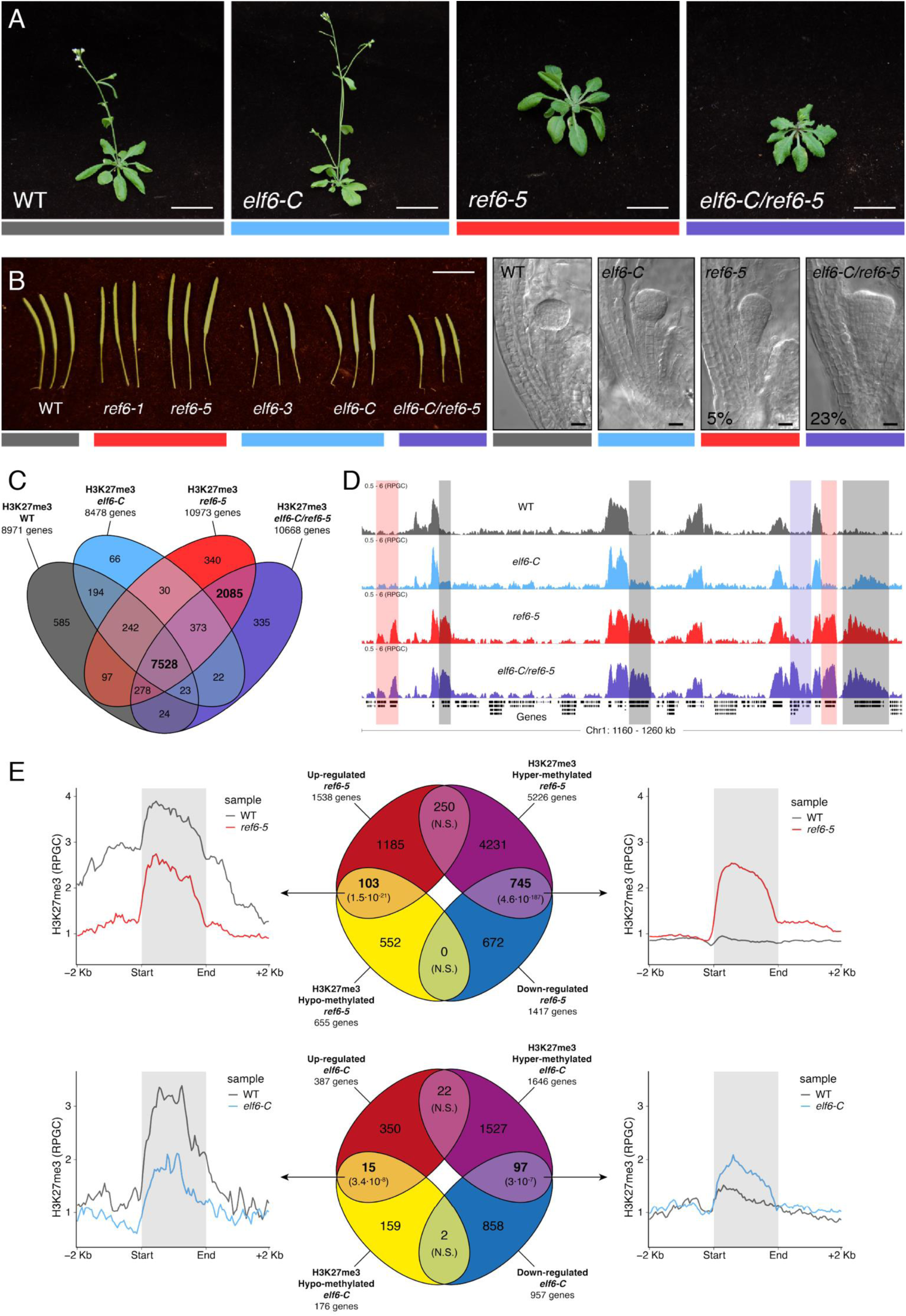
Arabidopsis Histone demethylases ELF6 and REF6 play distinct roles in development and H3K27me3 homeostasis. (A) Representative growth phenotypes of Arabidopsis wild-type (WT) and histone demethylase mutants (*elf6-C, ref6-5* and *elf6-C*/*ref6-5*). Scale bars, 1 cm. (B) Siliques and embryos from Arabidopsis wild-type (WT) and different mutant alleles of histone demethylase ELF6 and REF6. Numbers show the frequency of the abnormal embryos (n=250). Scale bars 1 cm and 10 μm, respectively. (C) Venn diagram showing the overlap between genes accumulating H3K27me3 in wild-type (WT) and histone demethylase mutants (*elf6-C, ref6-5* and *elf6-C*/*ref6-5*). (D) Genome browser views of background subtracted ChIP-seq signals as normalized reads per genomic content (RPGC). Shaded red boxes, genes targeted exclusively by REF6. Shaded grey boxes, genes targeted by REF6 and ELF6. Shaded purple boxes, genes genes targeted by both REF6 and ELF6, and only hyper-methylated in double mutant *elf6-C/ref6-5.* (E) Venn diagram showing overlap between differentially expressed genes (DEGs) and H3K27me3 differentially methylated genes in histone demethylase mutants. To the left metaplot for H3K27me3 levels for genes both up-regulated and hypo-methylated and to the right metaplot of H3K27me3 levels in genes both down-regulated and hyper-methylated. Top panel, *ref6-5*; Bottom panel, *elf6-C*. p-values for Fisher’s exact test are shown in brackets, N.S. Not Significant.

REF6 is thought to be a H3K27me3 demethylase and a positive regulator of gene expression (Hou, Zhou et al., 2014, Li, Gu et al., 2016, Lu et al., 2011, Wang, Gao et al., 2019), while the role of ELF6 remains poorly understood. To shed light on the function of these two proteins, we analyzed the distribution of H3K27me3 in *elf6-C, ref6-5* and *elf6-C/ref6-5* seedlings through ChIP-seq assays and compared the distribution of this epigenetic mark to that in wild-type plants. Overall, the accumulation of H3K27me3 at genes was more pronounced in *ref6-5* than in *elf6-C* (Fig 1C). Most of the hyper-methylated genes found in *elf6* (75%) were hyper-methylated to a greater extent in both *ref6-5* and *elf6-C/ref6-5*, suggesting that these histone demethylases have partially overlapping yet distinct roles in the control of H3K27me3 homeostasis in Arabidopsis (Fig. 1D and Supplemental Fig. S3-S4). In order to further understand the role of ELF6 and REF6 in transcriptional regulation, we performed an RNAseq analysis. When combining transcriptomic and H3K27me3 ChIP-seq data, we found a strong correlation primarily between genes that were both hyper-methylated at H3K27me3 and down-regulated, thus indicating that this epigenetic mark contributes to the transcriptional repression of these genes (Fig. 1E and Supplemental Fig.S5-S6). However, we also found genes that were hypo-methylated and up-regulated, which could be linked to the global transcriptional deregulation observed in these mutants. Taken together, our data point to the essential, yet distinct, roles of REF6 and ELF6 in H3K27me3 homeostasis at genic regions of the Arabidopsis genome.

### REF6 controls H3K27me1 homeostasis in chromatin

Biochemical analysis have revealed that REF6 and ELF6 can remove both tri- and di-methyl groups but not mono-methyl groups at lysine 27 on histone 3 (Lu et al., 2011). We therefore hypothesized that in addition to controlling H3K27me3 homeostasis, REF6 and ELF6 may be also implicated in H3K27me1 homeostasis. In order to test this hypothesis, we determined the distribution of H3K27me1 through ChIP-seq assays and found that most of the genes targeted by REF6 accumulate high levels of H3K27me1 in wild-type (Fig. 2A-C and Supplemental Fig. S7). Because the deposition of H3K27me1 in Arabidopsis is thought to be mediated by ATXR5 and 6 (Jacob et al., 2009, Jacob et al., 2010), we determined the genomic distribution of H3K27me1 in *atxr5/atxr6*. As previously described, H3K27me1 in these mutants was significantly reduced at pericentromeric heterochromatin but not affected in euchromatic regions (Supplemental Fig. S8). These data led us to hypothesize that the maintenance of H3K27me1 at euchromatin could be mediated by REF6. To validate this hypothesis, we investigated the relationship between H3K27me1 and H3K27me3 at genes targeted by REF6. This analysis revealed that the loss of REF6 activity results in both the accumulation of H3K27me3 and a drastic reduction of H3K27me1 at those loci, while the loss of ELF6 did not have an effect (Fig. 2B). We then assessed genomic regions directly targeted by REF6 (Cui et al., 2016, Li et al., 2016) and found that the accumulation of H3K27me3 in *ref6-5* was associated with a complete loss of H3K27me1 (Fig. 2C). Taken together these data revealed that the maintenance of H3K27me1 in euchromatin is dependent on REF6. Although it is well known that H3K27me1 in Arabidopsis contributes to the repression of heterochromatic TEs, its role in euchromatin remains unknown. To address this caveat, we examined the relationship between REF6-dependent H3K27me1 deposition and transcription. To aid this analysis, we divided the transcriptome into 10 quantiles of identical size according to their transcriptional state (Supplemental Fig. S9). We found that while H3K27me3 was primarily associated with strongly repressed genes in wild-type (first three quantiles), in *ref6-5*, the ectopic accumulation of H3K27me3 primarily affected genes that displayed low levels of expression (third to fifth quantiles) (Fig. 2D and Supplemental Fig. S10). Moreover, we found that the activity of REF6 was required by low-level expression genes (third to fifth quantile) (Fig. 2E and Supplemental Fig. S11). Collectively, these data support the view that REF6 contributes to both gene activation, by the removal of PRC2-dependent repressive marks, and to underpin low-level basal expression, by maintaining H3K27me1 in transcriptionally active chromatin.

**Figure 2.**
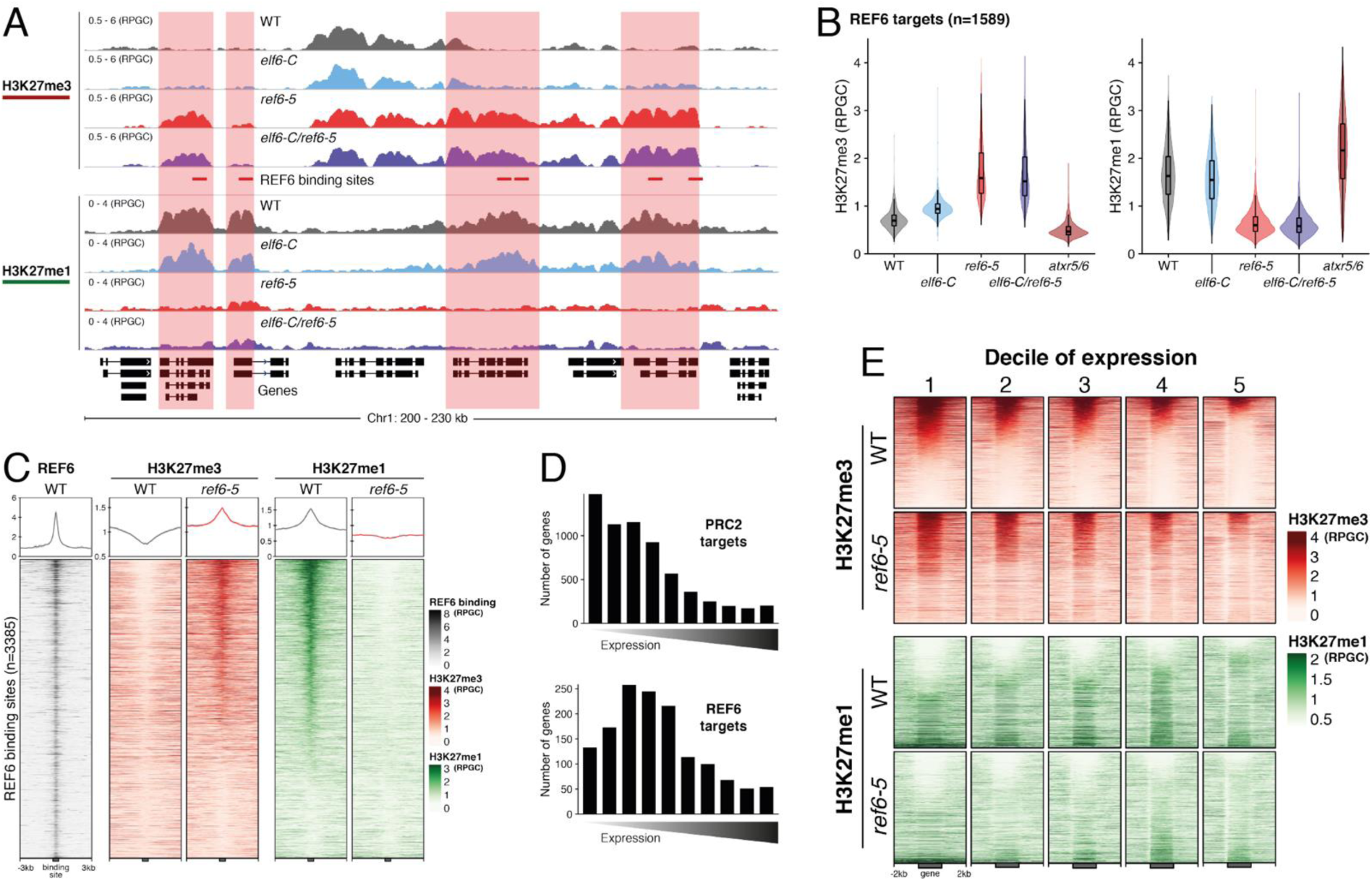
Arabidopsis REF6 play an essential roles in the deposition of H3K27me1 in active chromatin. (A) Genome browser views of background subtracted ChIP-seq signals for H3K27me3 and H3K27me1 as normalized reads per genomic content (RPGC) in wild-type (WT) and histone demethylase mutants (*elf6-C, ref6-5* and *elf6-C*/*ref6-5*). Shaded boxes, genes targeted exclusively by REF6. (B) Violin plots showing the distribution of H3K27me3 and H3K27me1 on genes targeted by REF6. Genes were categorised as targeted if a H3K27me3 peak was annotated on them in *ref6-5* and in *elf6-C*/*ref6-5* but not in WT. (C) Heatmap showing the distribution of H3K27me3 and H3K27me1 on genomic sequences targeted by REF6 for wild-type (WT) and *ref6-5* plants. Sample size n = 3,385. (D) Bar charts showing the number of genes for different expression quantiles predicted to be targeted by PRC2 and REF6. (E) Heatmap showing the distribution of H3K27me3 and H3K27me1 present on genes corresponding to low-expression (1-5) quantiles.

### Inheritance of ectopic H3K27me3 imprints alters the epigenome

It has been shown in Arabidopsis that histone demethylases are critical for the resetting of H3K27me3 across generations (Crevillén et al., 2014, Gan et al., 2014, Liu et al., 2019, Zheng et al., 2019). To understand the biological significance of this epigenetic resetting that likely takes place during plant sexual reproduction, we generated reciprocal crosses between *elf6-C/ref6-5* and wild-type plants. While F_1_ hybrids from these crosses were indistinguishable from the wild-type, a few F_2_ progenies (products of F_1_ self-pollination) displayed unexpected developmental phenotypes, including characteristics that were not present in either single or double mutants (n=1,500; 4.42% paternal transmission; 4.65% maternal transmission) (Fig 3A). Notably, some abnormal plants from F_2_ progenies were genetically wild-type for *ELF6* and *REF6*, and when we grew them we uncovered an array of developmental abnormalities that continued to segregate with stochastic frequencies in subsequent generations (Fig 3A). We reasoned that these phenotypes could have resulted from epimutations arising from the defective resetting of H3K27me3 during sexual reproduction in *elf6-C/ref6-5.* – a phenomenon not previously reported in plants. To test this hypothesis, we performed ChIP-seq analyses using seedlings from two independent epimutant F_5_ progeny. Our analysis revealed 535 euchromatic loci displaying elevated levels of H3K27me3, of which one third were also found to be hyper-methylated in the parental double mutant line used for reciprocal crosses (Fig. 3B). These data suggest that some of the H3K27me3 imprints present in epimutants were formed in *elf6-C/ref6-5* and were stably transmitted over five generations even after wild-type function was restored (Fig. 3C and Supplemental Fig. S12). We therefore named these lines *epiER* (<u>epi</u>mutants arising from *<u>elf6-C</u>/<u>ref6-5</u>)*. Notably, for both lines we found that the ectopic accumulation of H3K27me3 was particularly elevated in constitutive heterochromatin within the pericentromeric regions (Fig. 3D). Taken together, our data suggest that ELF6 and REF6 are necessary to limit the transmission of H3K27me3 imprints to offspring and that failure to do so results in epigenomic and developmental abnormalities.

**Figure 3.**
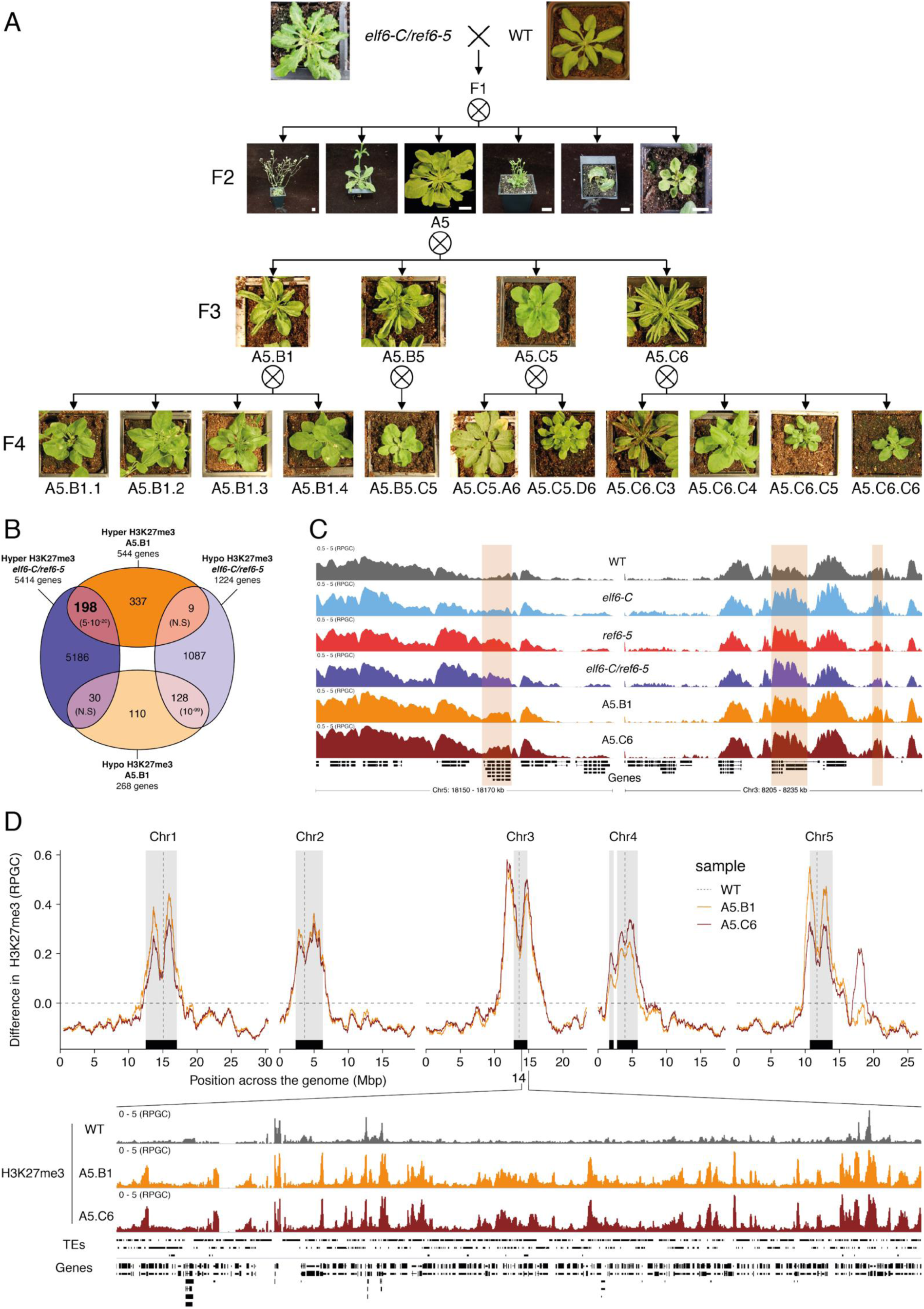
Pleiotropic developmental abnormalities associated with the inheritance of ectopic H3K27me3 imprints in Arabidopsis. (A) Abnormal phenotypes of plants from reciprocal crosses between *elf6-C/ref6-5* and wild-type (WT). Representative images of phenotypes arising from different progenies propagated by selfing. Bar chart, 1 cm. (B) Venn diagram showing the overlap in genes accumulating H3K27me3 in *elf6-C/ref6-5* and F_5_ progenies from A5.B1. p-values for Fisher’s exact test are shown in brackets, N.S. Not Significant. (C) Genome browser views of background subtracted ChIP-seq signals for H3K27me3 as normalized reads per genomic content (RPGC) in wild-type (WT), *elf6-C, ref6-5, elf6-C/ref6-5*, and F_5_ progenies from A5.B1 and A5.C6. Shaded boxes, genes showing transgenerational inheritance of H3K27me3. (D) Top panel: Differences in the chromosomal distribution of H3K27me3 as normalized reads per genomic content (RPGC) between F5 progenies from A5.B1 and A5.C6 and wild-type (WT). Grey shaded boxes, pericentromeric regions. Bottom panel: Genome browser view of ChIP-seq signal for H3K27me3 as normalized reads per genomic content (RPGC) in wild-type (WT), and F_5_ progenies from A5.B1 and A5.C6 in a pericentromeric region.

### Accumulation of ectopic H3K27me3 at centromeric heterochromatin is linked to DNA hypomethylation

Loss of DNA methylation has been linked to the abnormal deposition of H3K27me3 in heterochromatin (Batista & Kohler, 2020). However, mutants defective in H3K27me3 deposition do not affect global DNA methylation levels (Stroud, Do et al., 2014). To test if the ectopic accumulation of H3K27me3 found in *epiER*s could affect DNA methylation, we performed a BS-seq analysis on the two F_5_ epimutant progenies used for the ChIP-seq analysis. We found that both *epiER* lines displayed global reductions in DNA methylation, primarily at pericentromeric regions (Fig. 4A) (Miura, Yonebayashi et al., 2001, Vongs, Kakutani et al., 1993, Zemach, Kim et al., 2013). This global reduction in methylation occurred despite there being no ectopic accumulation of H3K27me3 in the parental mutant at any genes involved in the DNA methylation pathway. In addition, we found that the *epiERs* analysed displayed notable differences in DNA methylation levels between lines and among chromosomes (Fig 4A). In order to test if the observed differences and stochastic phenotypic segregation could be attributed to variation in DNA methylation between plants in the population, we performed a methylome analysis on individual plants. This analysis revealed that while some plants were consistently devoid of DNA methylation at pericentromeric regions, similar to the *ddm1* mutant, others displayed intermediate states that varied from chromosome to chromosome (Fig 4B). The loss of DNA methylation in constitutive heterochromatic regions is associated with a decrease in methylation at TEs and genes located therein (Fig. 4C and Supplemental Fig. S13). Since these pericentromeric regions fail to fully restore DNA methylation to wild-type levels and were elevated for H3K27me3 we hypothesized that they may be partially protected from the activity of the RNA-directed DNA methylation (RdDM) pathway, which establishes and maintains DNA methylation at euchromatic transposons and repetitive DNA elements in plants (Matzke & Mosher, 2014). To test this hypothesis, we investigated the relationship between DNA methylation and H3K27me3 on transposons located in euchromatic and constitutive heterochromatic genome regions. We found that in *epiERs*, heterochromatic TEs that gained H3K27me3 had a proportional loss of DNA methylation, whereas euchromatic TEs showed no change in DNA methylation (Fig. 4D). These data support the view that a gain in H3K27me3 has a negative effect on the deposition and/or the maintenance of DNA methylation at heterochromatic transposons. To evaluate the extent to which these defects may affect chromatin compaction, we performed immunostaining assays on interphase nuclei using specific antibodies. We found that in *epiERs* heterochomatin compaction is strongly affected (Supplemental Fig. S14) (Fig. 4E). Collectively, these data suggest that the ectopic accumulation of H3K27me3 in *epiERs* results in pericentromeric heterochromatin defects.

**Figure 4.**
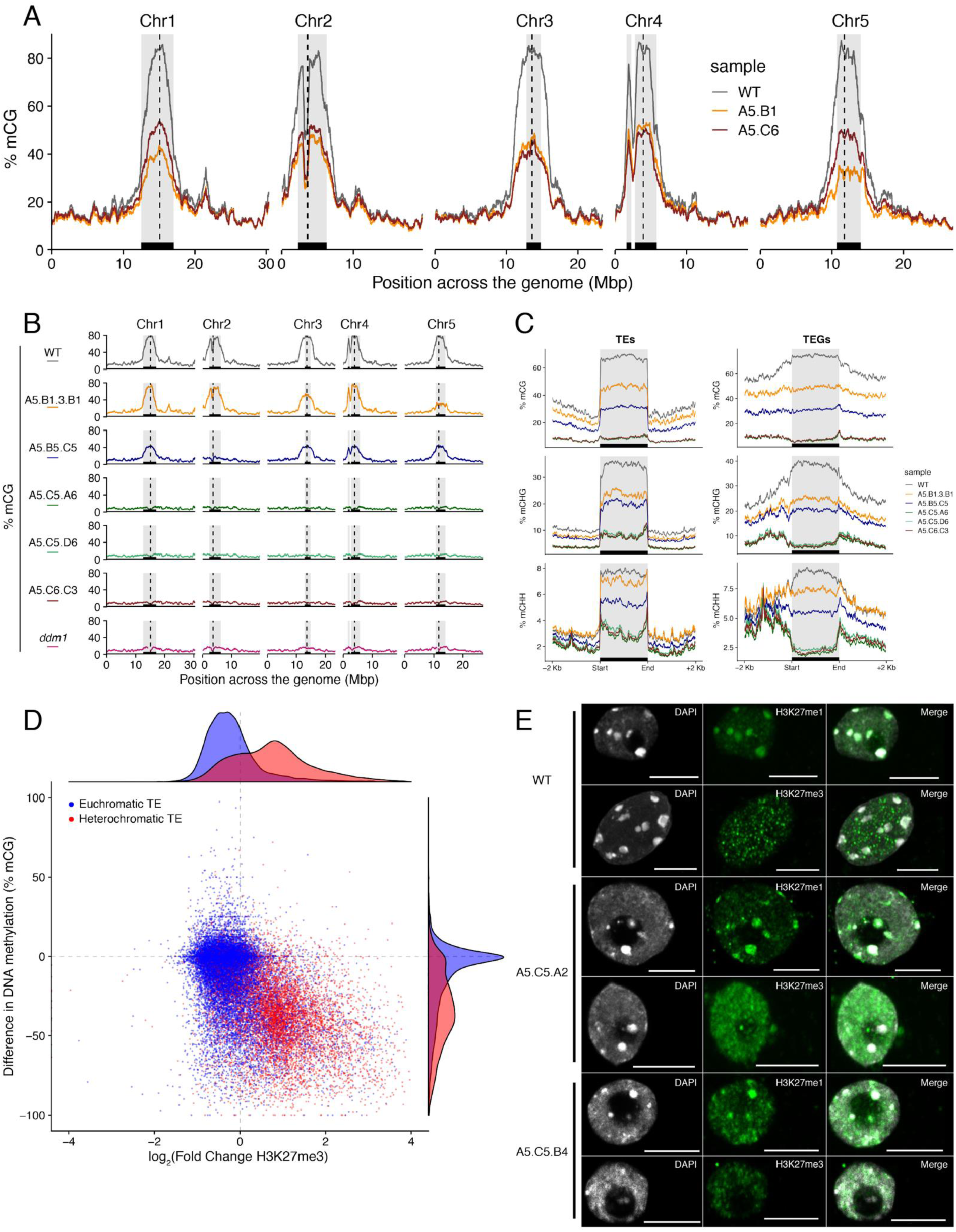
Ectopic accumulation of H3K27me3 is associated with the loss of DNA methylation at pericentromeric heterochromatin and affects chromatin condensation. (A) Distribution of DNA methylation across chromosomes of wild-type (WT) and progenies from *epiER*s A5.B1 and A5.C6. Grey shaded boxes, pericentromeric regions. (B) Distribution of DNA methylation across chromosomes of individual plants from wild-type (WT), *ddm1*, and *epiER*s A5.B1.3.B1, A5.B5.C5, A5.C5.A6, A5.C5.D6 and A5.C6.C3. Grey shaded boxes, pericentromeric regions. (C) Distribution of DNA methylation across Transposable Elements (TEs) and Transposable Element Genes (TEGs) of individual plants from wild-type (WT) and *epiER*s A5.B1.3.B1, A5.B5.C5, A5.C5.A6, A5.C5.D6 and A5.C6.C3. Black box, centromeric regions. (D) Correlation between DNA methylation changes and H3K27me3 changes on euchromatic and heterochromatic TEs, in wild-type (WT) and *epiER* A5.B1. (E) Immunolocalization showing the distribution of H3K27me3 and H3K27me1 in interphase nuclei of wild-type, A5.C5.A2 and A5.C5.B4 plants. Scale bars, 5 μm.

### Epigenomic defects result in transcriptional activation of pericentromeric loci and genome instability

We predicted that the abnormal distribution of epigenetic marks in ELF6/REF6-mediated epimutants could be responsible for the developmental abnormalities observed in these plants. To test this hypothesis, we performed a RNAseq analysis and found that 1,240 and 1,128 genes were misregulated in *epiER*s A5.B1 and A5.C6, respectively (Supplementary Table S1). A fraction of the upregulated in *epiER*s (483 and 544) were also upregulated in *elf6-C/ref6-5* plants (Fig 5A and Supplemental Fig S15A). Gene ontology analysis revealed that most upregulated genes in epimutants were involved in biotic stress responses (Fig 5B and Supplemental Fig S15B). When we investigated the chromosomal distribution of these deregulated genes, we found that some were located in constitutive pericentromeric heterochromatin and they showed the strongest upregulation effect (Fig 5C). These data suggest that the abnormal distribution of epigenetic marks in *epiERs* results in transcriptional activation of euchromatic and heterochromatic loci. Pericentromeric heterochromatin in plants is rich in TEs and is tightly regulated by DNA methylation and other epigenetic modifications (Dubin, Mittelsten Scheid et al., 2018), thus we hypothesized that the epigenomic perturbations found in *epiERs* could result in the activation of transposons. To test this hypothesis, we used our transcriptome data to determine the transcriptional state of different TEs in the two *epiER* progenies. We found that both RNA and DNA transposon families were significantly upregulated in *epiERs* (Fig 6A and Supplemental Fig S15). To assess whether the transcriptional activation of TEs in these epimutants could result in an increase in their mobility we determined their copy number in different *epiER* lines. We found that one heterochromatic transposon, CACTA1 (At2TE20205), and one euchromatic transposon, EVD (At5TE20395), showed a significant increase in copy number in both *epiERs* (Fig 6B). Further analysis revealed that these TEs were depleted in DNA methylation and significantly upregulated (Fig 6C and Supplemental FigS17). We then determined the precise location of some of the transposons newly mobilized in the different epimutants. We found that most novel insertions accumulated in euchromatin, continued to be active over multiple generations, and sometimes disrupted gene expression resulting in developmental phenotypes (Fig 6D-F and Supplemental Table S2). Collectively our data demonstrate that the developmental abnormalities found in *epiER* lines result from a combination of stably inherited genetic and epigenetic mutations.

**Figure 5.**
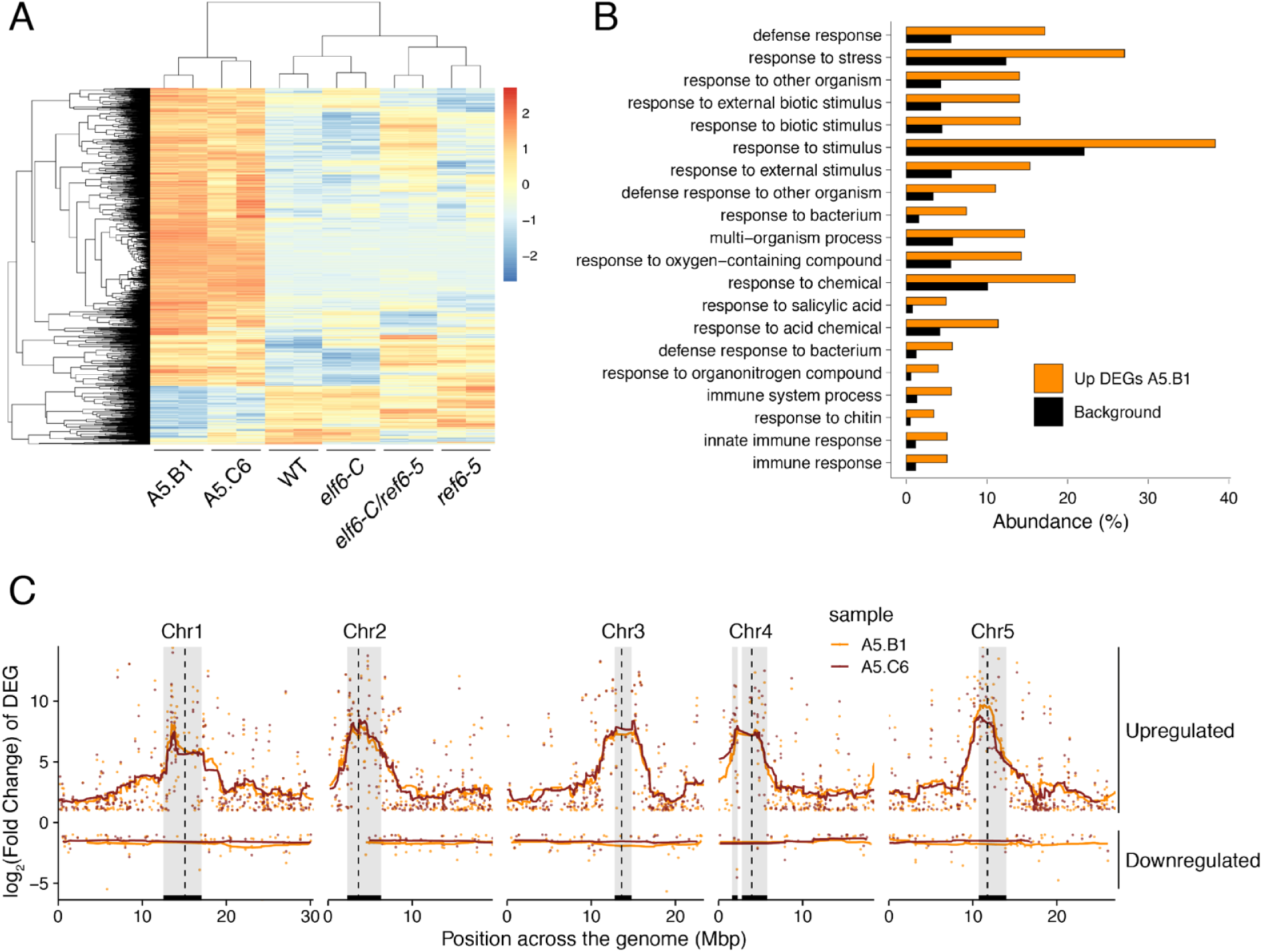
Global upregulation of centromeric gene expression in *epiER*s. (A) Heatmap showing scaled expression levels of Differentially Expressed Genes between wild-type and progeny of epi*ER* A5.B1 in wild-type (WT) *elf6-C, ref6-5, elf6-C/ref6-5*, and progenies of *epiER*s A5.B1 and A5.C6. (B) Gene Ontology analysis showing the functional categories enriched in genes upregulated in progeny of *epiER*s A5.B1. (C) Differential gene expression across each Arabidopsis chromosome for genes upregulated and downregulated in progenies of *epiER*s A5.B1 and A5.C6. Grey shaded boxes, pericentromeric regions

**Figure 6.**
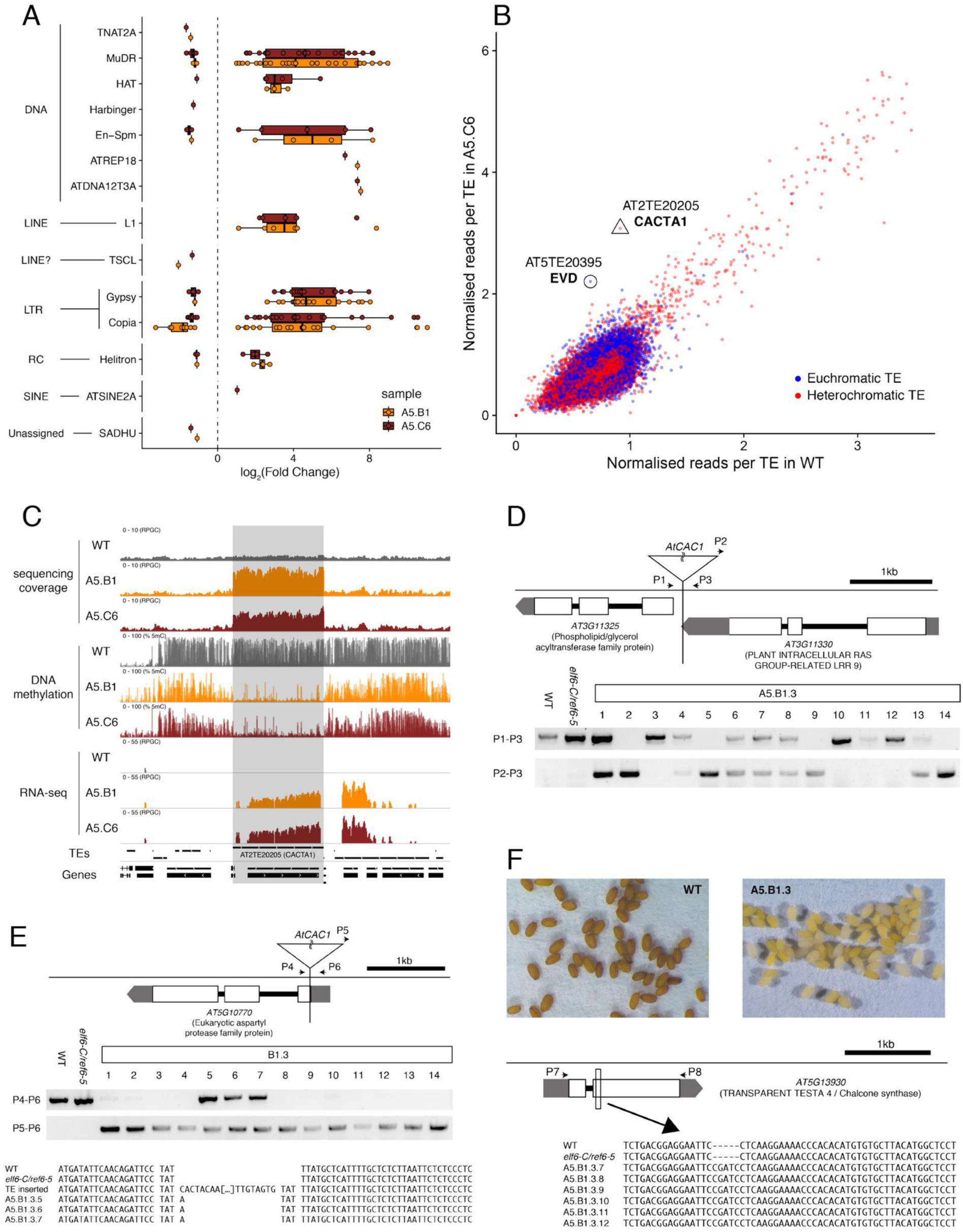
Transposon mobilization in *epiER*s results in heritable genetic lesions. (A) Differential expression of DNA and RNA transposon famikies grouped by superfamily in progenies of *epiER*s A5.B1 and A5.C6. (B) Copy number variation of transposons in progenies of *epiER* A5.C6. Blue dots, euchromatic TEs; Red dots, heterochromatic TEs. (C) Genome browser views of normalized sequencing coverage (RPGC), DNA methylation frequency (%) and RNAseq coverage (RPGC) in wild-type (WT) and progenies of *epiER*s A5.B1 and A5.C6. Grey box, AT2TE20205 (CACTA1). (D) Map of transposon insertion in AT3G11330 and its segregation in *epiER* A5.B1.3 progenies. P1-3, primers used for PCR amplification and sequencing. (E) Map of transposon insertion in AT5G10770 and sequence footprint resulting from re-mobilization in *epiER* A5.B1.3 progenies. P4-6, primers used for PCR amplification and sequencing. (F) Seed pigmentation defects caused by a sequence insertion in AT5G13930 (TRANSPARENT TESTA4/ CHALCONE SYNTHASE) resulting from transposon re-mobilization in in *epiER* A5.B1.3 progenies. P7-8, primers used for PCR amplification and sequencing.

## Discussion

In plants, histone modifications deposited by PRC2 play a critical role in growth and development, and in the adaptation of these processes to environmental fluctuations. Previous studies in Arabidopsis have shown that the activity of a distinct group of JmJ-type demethylases shape the genomic distribution of H3K27me3 (Yan et al., 2018). Three of these proteins – JMJ13, ELF6 and REF6 – have been shown to play important roles in development and in the regulation of environmental perception (Noh, Lee et al., 2004, Zheng et al., 2019). Our data show that REF6 and ELF6 regulate the removal of H3K27me3 at different genomic loci; while REF6 has a large repertoire of target genes, ELF6 activity is restricted to a small subset of genes, most of which can also be targeted by REF6. These data combined with our genetic analysis suggest that, despite the structural similarities between these two proteins, they play somewhat distinct functions in H3K27me3 homeostasis. Our data also support the view that although REF6 restricts the spreading of H3K27me3 to the genomic regions flanking PRC2 targets (Yan et al., 2018), it also plays an hitherto unrecognized role in the regulation of H3K27me1 homeostasis in euchromatin. This view is also supported by the overlap observed between REF6 genomic targets and the accumulation in wild-type plants of H3K27me1 in these regions, as well as by the complete loss of this chromatin mark in PRC2 target loci when REF6 activity is lost. Therefore, the deposition of H3K27me1 in Arabidopsis relies both on the activity of ATXR5 and ATXR6 in heterochromatin (Jacob et al., 2009, Jacob et al., 2010) and the activity of REF6 in transcriptionally active euchromatin (Supplemental FigS18). In mammals, the histone demethylases UTX and JMJD3, also known as KDM6A and KDM6B, have also been shown to catalyze the conversion of H3K27me3 and H3K27me2 into H3K27me1 (De Santa, Totaro et al., 2007, Lan, Bayliss et al., 2007, Lee, Villa et al., 2007, Swigut & Wysocka, 2007). Moreover, defects in PRC2 methyltransferase activity in mammals completely abolish the accumulation of H3K27me1 in embryonic stem cells (Ferrari, Scelfo et al., 2014, Montgomery, Yee et al., 2005), suggesting a conserved PRC2-mediated mechanism for H3K27me1 homeostasis in euchromatin, in both animals and plants. The precise mechanism responsible for the deposition of this chromatin mark in Arabidopsis is currently unknown, but two possible scenarios are envisaged: either the deposition of H3K27me1 in euchromatin is dependent on the activity of PRC2 and REF6, or this mark could be deposited by an unknown histone mono-methyltransferase which requires REF6 for its maintenance (Fig. 7). In mammals the presence of H3K27me1 in actively transcribed genome regions has been associated with the promotion of transcription (Ferrari et al., 2014). This may explain why, in Arabidopsis, genes associated with H3K27me1 display moderate levels of expression whereas the conversion of this mark into H3K27me3 negatively impacts their transcriptional rate. Plant somatic cells accumulate H3K27me3 primarily at protein-coding genes, however, in reproductive tissues and mutants where DNA methylation is reduced, this mark also accumulates at transposon loci (Deleris, Stroud et al., 2012, Weinhofer, Hehenberger et al., 2010). Other studies have also reported the accumulation of H3K27me3 in transposon sites for plant species with reduced levels of DNA methylation (Montgomery et al., 2005), as well as in mammal somatic and reproductive tissues which also show a reduction in DNA methylation (Hanna, Perez-Palacios et al., 2019, Reddington, Sproul et al., 2014, Saksouk, Barth et al., 2014). However, our data does not fully support the idea that the deposition of this chromatin mark acts as a compensatory system to silence hypomethylated TEs (Deleris et al., 2012, Hanna et al., 2019). Instead, our results suggest that the homeostasis and function of H3K27me1 and H3K27me3 in plants is more complex than previously anticipated.

**Figure 7.**
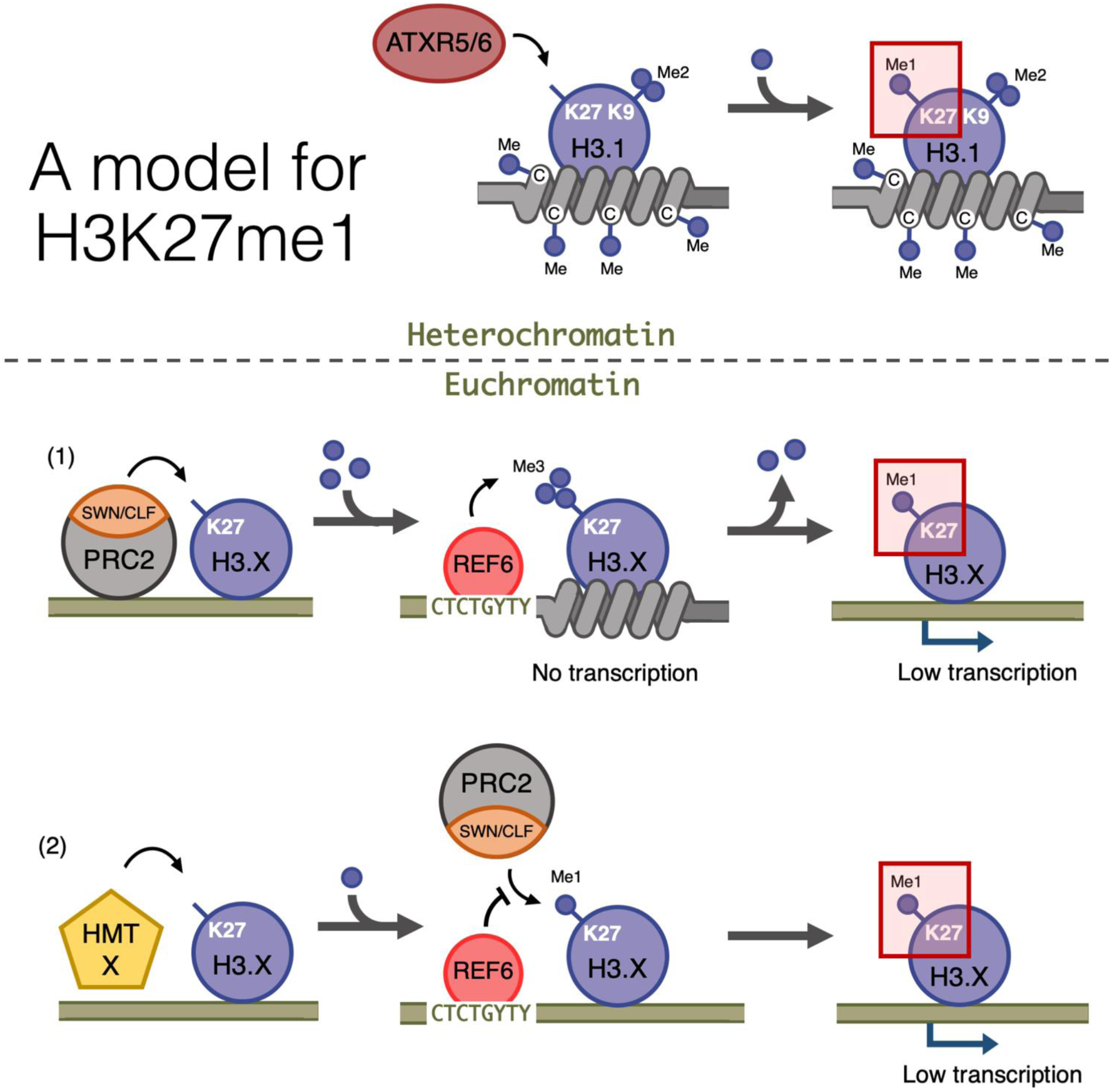
Proposed model for the deposition of H3K27me3 and H3K27me1 in Arabidopsis. Hypothetical model for the distinct mechanisms of deposition of H3K27me3 and H3K27me1 in different chromatin compartments. In pericentromeric heterochromatin, ATRX5/6 deposits in one one-step H3K27me1 in histones accumulating H3K9me2. In euchromatin, the H3K27me3 deposited by histone methylases SWN/CLF from the PRC2 complex is converted to H3K27me1 by the activity of histone demethylase REF6. Alternatively, the deposition of H3K27me1 is mediated in one one-step process by a yet unknown histone methylase and maintained by the activity of REF6 to prevent the deposition of additional methyl groups by PRC2. Dark blue circles, methyl groups. C, represent Cytosines. H3.X represents a H3 variant that is not H3.1. Coiled lines represent closed and inactive chromatin.

The stable inheritance of *de novo* acquired DNA methylation imprints in plants are well documented. Mutations in the machinery involved in the deposition of DNA methylation, such as the cytosine *DNA METHYLTRANSFERASE 1* (*MET1)* and the chromatin-remodeling *DEFICIENT IN DNA METHYLATION 1* (DDM1), lead to the formation of epimutations caused by DNA hypomethylation (Johannes, Porcher et al., 2009, Kakutani, Jeddeloh et al., 1996, Mathieu, Reinders et al., 2007). These epimutations are maintained during sexual reproduction and remain stable over several generations, even after the function of *MET1* or *DDM1* is restored. Moreover, natural epimutations created during asexual propagation and associated with DNA hypomethylation are associated with TEs and can be stable over multiple generations thus contributing to novel yet stable phenotypic variation (Ong-Abdullah, Ordway et al., 2015, Wibowo, Becker et al., 2018). Despite accumulating evidence for the active role of histone demethylases in resetting H3K27me3 at specific loci during sexual reproduction (Crevillén et al., 2014, Noh et al., 2004), the precise mechanism(s) remain unknown. Our data show that a failure to reset H3K27me3 during sexual reproduction results in the trans-generational inheritance of this chromatin mark in euchromatin, even when functional demethylase activity is restored. One possible explanation for these findings could be that some of the H3K27me3 imprints that are ectopically deposited in histone demethylase mutants cannot be reset because they are distal to the target sequences recognized by these demethylases. Once established, these H3K27me3 imprints could be maintained across generations as epimutations, which we termed *epiER*, through the recruitment of LHP1-PRC2 complexes (Derkacheva, Steinbach et al., 2013). Our data also revealed that the inheritance of these imprints causes defects in the maintenance of DNA methylation at heterochromatic regions of the genome. The ectopic deposition of H3K27me3 in constitutive heterochromatin may be linked to defects in the resetting of DNA methylation thought to take place during gametogenesis (Calarco, Borges et al., 2012, Ibarra, Feng et al., 2012, Slotkin, Vaughn et al., 2009) and/or early embryo development (Bouyer, Kramdi et al., 2017). Under this scenario, an active resetting of H3K27me3 in gametes would be critical for the re-establishment of DNA methylation in heterochromatin after fertilization. Moreover, epigenomic alterations could explain the heritable, yet unstable, phenotypes observed in *epiER*s. Similar epimutations and phenotypic variation have been shown to arise from crosses between wild-type plants and mutants defective in the machinery that maintain DNA methylation (Kakutani et al., 1996, Kato, Takashima et al., 2004, Marí-Ordóñez, Marchais et al., 2013, Mirouze, Lieberman-Lazarovich et al., 2012). As in these studies, we also found that *epiER*s have defects in the silencing of some transposons resulting in an increase in genetic lesions associated with their mobilization.

Taken together our data reveal novel, critical roles for histone demethylases in maintaining both genome integrity and transcriptional states during plant development.

## Acknowledgments

We thank Gary Grant for help with plant husbandry; Xiaofeng Cao and Caroline Dean for seed stocks and data. Ranjth Papareddy for the identification of *ref6-5* and Liliana M. Costa for discussions and comments on the manuscript. Supported by the ERC AdG IMMUNEMESIS Project, the DFG SPP1529 Program, and the Max Planck Society (D.W.), ANR/CNRS grant (EpiGEN) to M.B., JSPS grant (JP19H05676) to

M.U. and BBSRC grants (BB/L003023/1, BB/N005279/1, BB/N00194X/1 and BB/P02601X/1) to J.G-M.

## Author contribution

M.N., J.A.S., M.B. and JG-M conceived the project. M.N., S.O., D.L., C.R., Y.H., J.S.R-P., F.A., A.W., M.U. and JG-M designed and conducted experiments. M.N., J.A.S., A.D., S.O., D.L., D.M., J.D., M.U., M.B. and JG-M analysed the data. M.B. and JG-M wrote the manuscript with input from the rest of the authors.

## Data and Materials Availability

Sequence data (BS-seq, RNA-seq and ChiP-seq) that support the findings of this study have been deposited at the European Nucleotide Archive (ENA) under the accession code PRJEB36508.

---

**Supplemental Figure S1. Graphical representation of mutant alleles for ELF6 and REF6 used in this study.**

(A) Schematic diagram of *ELF6* locus showing the location of different mutant alleles: *elf6*-3 is a T-DNA insertion (SALK_074694) in the first exon and *elf6-C* is a CRISPR/Cas9 deletion in the first exon that leads to an early stop codon. Black boxes represent exons, triangle shows the insertion site for *elf6*-3 and red arrows mark the start and end of the deleted region in *elf6-C*. Scale bar, 1 kb.

(B) Schematic diagram of *REF6* locus showing the location of different mutant alleles: *ref6-5* (GABI_70E03) and *ref6*-1 (SALK_001018) are T-DNA insertions in the sixth and last exon, respectively. Black boxes represent exons and triangles shows the T-DNA insertion sites. Scale bar, 1 kb.

**Supplemental Figure S2. Phenotypic characterization of *elf6-C, ref6-5* and double mutants.**

(A) Rossete leaves at bolting stage for Arabidopsis wild-type (WT) and histone demethylase mutants (*elf6-C, ref6-5* and *elf6-C*/*ref6-5*). Scale bars, 1 cm.

(B) Boxplot for leaf number at bolting stage for wild-type (WT) and different mutants. Letters represent groups of statistically significantly different samples. Differences between genotypes determined by Students t-test, using a sample size of n = 30.

(C) Growth phenotypes of wild-type (WT) and *elf6-C*/*ref6-5* mutant. Scale bar, 1 cm.

(D) Boxplot of plant height for wild-type (WT) and histone demethylase mutants. Asterisks represent significant differences between samples. Differences between genotypes determined by Students t-test test, **** p<0.001, sample size of n = 30.

(E) Boxplot of seed germination rates in wild-type (WT) and histone demethylase mutants. Germination was scored as radicle protrusion through seed coat. n = 300 from six biological replicates.

**Supplemental Figure S3. Genes with differential K3K27me3 methylation in histone demethylase mutants.**

(A) Euler diagram of H3K27me3 hyper-methylated genes in histone demethylase mutants compared to WT.

(B) Euler diagram of H3K27me3 hypo-methylated genes in histone demethylase mutants compared to WT.

**Supplemental Figure S4. Genomic regions accumulating K3K27me3 in histone demethylase mutants.**

(A) Genome browser view of ChIP-seq signal as normalized reads per genomic content (RPGC). Shaded red boxes, genes targeted by REF6.

(B) Genome browser view of ChIP-seq signal as normalized reads per genomic content (RPGC). Shaded blue box, gene targeted exclusively by ELF6.

(C) Genome browser view of ChIP-seq signal as normalized reads per genomic content (RPGC). Shaded purple boxes, genes targeted by both REF6 and ELF6, and only hyper-methylated in the double mutant *elf6-C*/*ref6-5*.

**Supplemental Figure S5. Genes differentially expressed in histone demethylase mutants.**

(A) Euler diagram of down-regulated genes in histone demethylase mutants compared to WT.

(B) Euler diagram of up-regulated genes in histone demethylase mutants compared to WT.

**Supplemental Figure S6. Relation between hypermethylation of downregulation in *elf6-C*/*ref6-5*.**

Venn diagram showing overlap between differentially expressed genes (DEGs) and H3K27me3 differentially methylated genes in *elf6-C*/*ref6-5*. Venn diagram showing overlap between differentially expressed genes (DEGs) and H3K27me3 differentially methylated genes in *elf6-C*/*ref6-5*. To the left metaplot for H3K27me3 levels for genes both up-regulated and hypo-methylated and to the right metaplot of H3K27me3 levels in genes both down-regulated and hyper-methylated. p-values for Fisher’s exact test are shown in brackets, N.S. Not Significant.

**Supplemental Figure S7. REF6 catalyses H3K27me3 to H3K27me1 in genes causing derepression.**

(A) Heatmap showing the distribution of H3K27me3 (red) and H3K27me1 (green) on genes targeted by REF6. Genes were categorised as targeted if a H3K27me3 peak was annotated on them in *ref6-5* and in *elf6-C*/*ref6-5* but not in WT. n=1589. Intensity of the colour represents RPGC of ChIP-seq. Genes are sorted by the amount of H3K27me3 in WT. Boxes at the top represent metaplots of the average signal for all these genes.

(B) Metaplot of median of H3K27me3 and H3K27me1 across both downregulated and hypermethylated genes in *elf6-C*/*ref6-5* for wild-type (WT) and histone demethylase mutants, n=968.

(C) Heatmap of the same genes as Fig S 7B. Intensity of the colour represents ChIP-seq signal as RPGC.

(D) Metaplot of median of H3K27me3 and H3K27me1 across both upregulated and hypomethylated genes in *elf6-C*/*ref6-5* for wild-type (WT) and histone demethylase mutants, n=256.

(E) Heatmap of the same genes as Fig S 7D. Intensity of the colour represents ChIP-seq signal as RPGC.

**Supplemental Figure S8. ATXR5/6 contributes to the deposition of H3K27me1 in pericentromeric regions.**

(A) Distribution of H3K27me1 across chromosomes of wild-type (WT) and *atxr5/6* mutants. Grey shaded boxes, pericentromeric regions.

(B) Distribution of H3K27me1 across Chromosome 4 of Arabidopsis genome in wild-type (WT) and *atxr5/6* mutants. Grey shaded boxes, pericentromeric regions.

**Supplemental Figure S9. Division of Arabidopsis genes according to their levels of expression.**

Violin plot with boxplots representing the expression levels of genes in Arabidopsis divided in 10 quantiles.

**Supplemental Figure S10. Histone demethylase mutants show an increase in H3K27me3 on genes with intermediate levels of expression**

Heatmaps showing the H3K27me3 ChIP-seq signal across genes split by 10 quantiles of expression for wild-type (WT) and histone demethylase mutants. Boxes on top represent metaplots of the median signal in each quantile. Genes are sorted by the amount of H3K27me3 in WT. (A) WT. (B) *elf6-C*. (C) *ref6-5*. (D) *elf6-C*/*ref6-5*.

**Supplemental Figure S11. Distribution of different epigenetic features for different expression quantiles.**

Heatmaps showing signal for different epigenetic features across 10 expression quantiles for wild-type. Boxes on top represent metaplots of the median signal in each quantile. Genes in all panels are always sorted by the amount of H3K27me3. (A) H3K27me3. (B) H3K27me1. (C) H3K9ac. (D) ATAC-seq. (E) PolII-seq. (F) Mnase-seq.

**Supplemental Figure S12. Inheritance of ectopic H3K27me3 in *epiERs*.**

(A) Venn diagram showing the intersection between genes showing accumulation of H3K27me3 in *elf6-C*/*ref6-5* and *epiER* A5.C6. p-values for Fisher’s exact test are shown in brackets, N.S. Not Significant.

(B) Metaplot of the median of ChIP-seq RPGC across genes hypermethylated in both *elf6-C*/*ref6-5* and *epiER* A5.C6. n=198.

(C) Euler diagram showing the intersection between the genes hypermethylated in *elf6-C*/*ref6-5* and *epiERs* A5.B1 and A5.C6.

**Supplemental Figure S13. DNA hypomethylation of transposable elements in *epiERs*.**

Metaplot showing the proportion of DNA methylation across transposable elements (A) and transposable element genes (B) for wild-type (WT) and progenies from two *epiERs*.

**Supplemental Figure S14. Condensation of chromatin in *epiERs*.**

Bar plot showing the fraction of nuclei categorised as decondensed after DAPI staining in wild-type (WT) and *epiERs* A5.C5.A2, A5.C5.B4 and A5.C5.C1. Number of nuclei for each category in each line shown inside plot.

**Supplemental Figure S15. Genes upregulated in *epiERs*.**

(A) Heatmap showing scaled expression levels of Differentially Expressed Genes between wild-type and progeny of epi*ER* A5.C6 in wild-type (WT) *elf6-C, ref6-5, elf6-C/ref6-5*, and progenies of *epiER*s A5.B1 and A5.C6..

(B) Gene Ontology analysis showing the functional categories enriched in genes upregulated in *epiER* A5.C6.

**Supplemental Figure S16. Transcriptional upregulation of transposons in *epiERs*.**

Heatmap showing scaled logged expression of all the families of transposons or Arabidopsis. Samples represented are wild-type (WT), histone demethylase mutants and *epiERs* A5.B1 and A5.C6.

**Supplemental Figure S17. Deregulation of EVD transposon in *epiERs*.**

Genome browser view showing normalised sequencing coverage (RPGC), DNA methylation rate (%) and RNA-seq (RPGC) in wild-type (WT) and progenies from *epiERs* A5.B1 and A5.C6. Grey box, AT2TE20395 (EVD).

## Methods

### Plant material and Plant Growth

All plant lines used in this study were derived from *Arabidopsis thaliana* Col-0 accession. The T-DNA insertion lines *ref6-1* (SALK_001018), *elf6-3* (SALK_074694), *atxr5* (SALK_130607) and *atxr6* (SAIL_240_H01) have been previously described. The *ref6-5* mutant (GABI_705E03) was obtained from the GABI-Kat collection (Kleinboelting, Huep et al., 2012). The genomic deletion in *elf6-C* was produced using two sgRNAs (Table S1) and CRISPR/Cas9 (Durr, Papareddy et al., 2018). Double mutants were produced by hand crossing. The plant materials used for crossing and flowering time measurements were grown in chambers under long day conditions (16 h light, 8 h dark) with 120 μ mol m^-2^ s^-1^ light intensity (22°C daytime, 20°C at night). Plants for the screening were grown in a climate-controlled greenhouse under long day conditions (20°C daytime, 20°C at night,16 h light plus 8 h dark). The seeds were mixed in 0.1% Agarose and underwent 2 d cold treatment at 4°C in the dark. After treatment seeds were directly sown on soil and transferred to growth a chamber or greenhouse.

### Genotyping and Phenotyping

Primary transformants were identified using the seed-specific RFP reporter under a Leica MZ-FL III stereomicroscope (Leica Camera AG). Genotyping of CRISPR/Cas9-based mutations and T-DNA insertions were performed using KAPA-Taq (Sigma-Aldrich) following the manufacturer’s instructions. PCR product size was selected using gel electrophoresis and the introduced genetic lesion was determined by sequencing (See Supplementary Table 3 and Supplementary Fig. S1). The phenotypes of whole plants, leaf number and rosette size were scored at bolting. Silique length measurement were carried out on the 6^th^-15^th^ siliques of main stems, when the last flowers of the inflorescence started producing siliques. The mean value of the 10 siliques represented the silique length of a plant. For embryo analysis, ovules from self-pollinated plants were cleared with a chloral hydrate solution, observed with a light microscope (Zeiss AxioImager A2) and photographed with a digital camera (Zeiss AxioCam HRm).

### ChIP-seq assay

ChIP-seq assays were performed on 14 days old *in vitro* shoot seedlings using anti-H3K27me3 (Millipore 07-449) or anti-H3K27me1 (Millipore 07-448), following a procedure modified from Gendrel, Lippman et al. (2005). Five grams of plantlets were cross-linked in 1% (v/v) formaldehyde at room temperature for 15mn. Crosslinking was then quenched with 0.125 M glycine for 5 min. The crosslinked plantlets were ground and nuclei were isolated and lysed in Nuclei Lysis Buffer (1% SDS, 50mM Tris-HCl pH 8, 10mM EDTA pH 8). Cross-linked chromatin was sonicated using a water bath Bioruptor UCD-200 (Diagenode, Liège, Belgium) (15s on/15s off pulses; 15 times). The complexes were immunoprecipitated with antibodies, overnight at 4°C with gentle shaking, and incubated for 1 h at 4°C with 40 µL of Protein AG UltraLink Resin (Thermo Scientific). The beads were washed 2 × 5 min in ChIP Wash Buffer 1 (0.1% SDS, 1% Triton X-100, 20 mM Tris-HCl pH 8, 2 mM EDTA pH 8, 150 mM NaCl),2 × 5 min in ChIP Wash Buffer 2 (0.1% SDS, 1% Triton X-100, 20 mM Tris-HCl pH 8, 2 mM EDTA pH 8, 500 mM NaCl), 2 × 5 min in ChIP Wash Buffer 3 (0.25 M LiCl, 1% NP-40, 1% sodium deoxycholate,10 mM Tris-HCl pH 8, 1 mM EDTA pH 8) and twice in TE (10 mM Tris-HCl pH 8, 1 mM EDTA pH 8).ChIPed DNA was eluted by two 15-min incubations at 65°C with 250 μL Elution Buffer (1% SDS, 0.1 M NaHCO_3_). Chromatin was reverse-crosslinked by adding 20 μL of NaCl 5M and incubated over-night at 65°C. Reverse-cross-linked DNA was submitted to RNase and proteinase K digestion, and extracted with phenol-chloroform. DNA was ethanol precipitated in the presence of 20 μg of glycogen and resuspended in 50 μL of nuclease-free water (Ambion) in a DNA low-bind tube. 10 ng of IP or input DNA was used for ChIP-Seq library construction using NEBNext® Ultra DNA Library Prep Kit for Illumina® (New England Biolabs) according to manufacturer’s recommendations. For all libraries, 12 cycles of PCR were used. The quality of the libraries was assessed with Agilent 2100 Bioanalyzer (Agilent).

### Computational analysis of ChIP-seq

Single-end sequencing of ChIP samples was performed using Illumina NextSeq 500 with a read length of 76 bp. Reads were quality controlled using FASTQC (http://www.bioinformatics.babraham.ac.uk/projects/fastqc/). Trimmomatic was used for quality trimming. Parameters for read quality filtering were set as follows: Minimum length of 36 bp; Mean Phred quality score greater than 30; Leading and trailing bases removal with base quality below 5. The reads were mapped onto the TAIR10 assembly using Bowtie (Langmead, 2010) with mismatch permission of 1 bp. To identify significantly enriched regions, we used MACS2 (Zhang, Liu et al., 2008). Parameters for peaks detection were set as follows: Number of duplicate reads at a location:1; mfold of 5:50; q-value cutoff:0.05; extsize 200; broad peak. Visualization and analysis of genome-wide enrichment profiles were done with IGB. Peak annotations such as proximity to genes and overlap on genomic features such as transposons and genes were performed using BEDTOOLS INTERSECT. To identify regions that were differentially enriched in the H3K27me3 or H3K27me1 histone modification between WT and mutants, we used DIFFREPS (Shen, Shao et al., 2013) with parameters of pvalue 0,05; z-score cutoff 2; windows 1000.

### Expression profiling by RNA-seq

Leaf samples were collected from 4 wk old plants. Total RNA was extracted using RNeasy Plant Mini Kit (Qiagen) according to manufacturer’s instructions and used to produce libraries using TruSeq RNA library Prep Kit v2 (Illumina). Pooled libraries were sequenced in a NextSeq®550 sequencing platform (Illumina). Two biological replicates were generated for each genotype, and at least 20 million reads were produced per replicate.

### Generation of epimutations using histone demethylase mutants

The second generation of homozygous *elf6-C*/*ref6-5* were crossed reciprocally to wild-type plants (Col-0). F_1_ progenies were self-pollinated to generate F_2_ seeds that were grown in individual pots until bolting and the frequency of developmental phenotypes was scored. Plants displayed developmental phenotypes not found in *elf6-C, ref6-5 or elf6-C*/*ref6-5* mutants where genotyped by PCR to determine their zygosity.

### Bisulfite sequencing

Rosette leaves from five plants were pooled for each sample. Genomic DNA was extracted with the DNeasy Plant Mini Kit (Qiagen, Germany). DNA libraries were generated using the Illumina TruSeq Nano kit (Illumina, CA, USA). DNA was sheared to 350 bp. The bisulfite treatment step using the Epitect Plus DNA Bisulfite Conversion Kit (Qiagen, Germany) was inserted after the adaptor ligation; incubation in the thermal cycler was repeated once before clean-up. After clean-up of the bisulfite conversion reaction, library enrichment was done using Kapa Hifi Uracil+ DNA polymerase (Kapa Biosystems, USA). Libraries were sequenced with 2 x 150 bp paired-end reads on an HiSeq 4000 (Illumina), with conventional gDNA libraries in control lanes for base calling calibration. Sixteen to twenty four libraries with different indexing adapters were pooled in each lane.

### Computational analysis of paired end BS-seq

Paired-end quality was assessed using FASTQC (Andrews, Krueger et al., 2010). Trimmomatic (Bolger, Lohse et al., 2014) was used for quality trimming. Parameters for read quality filtering were set as follows: Minimum length of 40 bp; sliding window trimming of 4 bp with required Phred quality score of 20. Trimmed reads were mapped to the *Arabidopsis thaliana* TAIR10 genome assembly using bwa-meth (Pedersen, Eyring et al., 2014) with default parameters. Mapped reads were deduplicated using picardtools (Picard toolkit, 2019), and numbers of methylated/unmethylated reads per position were retrieved using MehtylExtract (Oliver, Barturen et al., 2014) and custom scripts.

### Pericentromeric heterochromatic regions

Heterochromatin regions were defined as in Qiu, Mei et al. (2019)(Chr1:12,500,000– 17,050,000, Chr2:2,300,000–6,300,000, Chr3: 12,800,000–14,800,000, Chr4: 1,620,000–2,280,000; 2,780,000–5,804,000, Chr5: 10,680,000–14,000,000).

### Gene expression and ontology analysis

We used agriGO v2.0 (Tian, Liu et al., 2017) to classify significantly enriched Gene Ontology (GO) terms associated with differential expression.

### Immunostaining of chromatin

Leaf protoplasts were isolated and fixed. After rehydration in PBS, slides were blocked in 2% BSA in PBS (30 min, 37°C) and incubated overnight at 4°C in 1% BSA in PBS containing antibodies (Upstate Biotechnology) specific to lysine-27-monomethylated H3 (1:100 dilution), and lysine-27-trimethylated H3 (1:100 dilution). Detection was carried out with an FITC-coupled antibody to rabbit IgG (Molecular Probes; 1:100 dilution, 37°C, 40 min) in 0.5% BSA in PBS. DNA was counterstained with 4,6 diamidino-2-phenylindole (DAPI) in Vectashield (Vector Laboratories).

### Data visualisation

For visualising BS-seq, RNA-seq and ChIP-seq genomic data we used Integratice Genomic Viewer (IGV) (Thorvaldsdóttir, Robinson et al., 2013), And R version 3.5.1 (www.r-project.org) with packages ggplot2 (Wickham, 2016), eulerr (Larsson, 2019), pheatmap (Kolde, 2015) and EnrichedHeatmap (Gu, Eils et al., 2018).

### Prediction of new TE insertion sites and molecular validation

We analysed Bisulfite-seq data using Bismark (Krueger & Andrews, 2011) using the following parameters:–bowtie2 –ambiguous –unmapped –R 10 –score_min L,0,-0.6 - N 1. Identification of new TE insertion sites was performed using epiTEome (Daron & Slotkin, 2017). For the validation of new transposon insertions, we designed primers outside of predicted TE insertion site and inside the transposon based on physical reads identified by epiTEome. We used KAPA Taq Polymerase and PCR conditions of 95°C for 5 min, followed by 30-35 cycles of 95°C for 30 s, 58°C for 15 s, and 72°C for 2 min. The list of primers employed for this analysis are listed (Supplementary Table S1).

## References

Andrews S, Krueger F, Segonds-Pichon A, Biggins L, Krueger C, Wingett S (2010) FastQC: a quality control tool for high throughput sequence data. In Babraham, UK: Babraham Institute

Batista RA, Kohler C (2020) Genomic imprinting in plants-revisiting existing models. Genes Dev 34: 24–36

Berger SL (2007) The complex language of chromatin regulation during transcription. Nature 447: 407–12

Bolger AM, Lohse M, Usadel B (2014) Trimmomatic: a flexible trimmer for Illumina sequence data. Bioinformatics 30: 2114–2120

Bouyer D, Kramdi A, Kassam M, Heese M, Schnittger A, Roudier F, Colot V (2017) DNA methylation dynamics during early plant life. Genome Biol 18: 179

Calarco JP, Borges F, Donoghue MT, Van Ex F, Jullien PE, Lopes T, Gardner R, Berger F, Feijo JA, Becker JD, Martienssen RA (2012) Reprogramming of DNA methylation in pollen guides epigenetic inheritance via small RNA. Cell 151: 194–205

Crevillén P, Yang H, Cui X, Greeff C, Trick M, Qiu Q, Cao X, Dean C (2014) Epigenetic reprogramming that prevents transgenerational inheritance of the vernalized state. Nature 515: 587–590

Cui X, Lu F, Qiu Q, Zhou B, Gu L, Zhang S, Kang Y, Cui X, Ma X, Yao Q, Ma J, Zhang X, Cao X (2016) REF6 recognizes a specific DNA sequence to demethylate H3K27me3 and regulate organ boundary formation in Arabidopsis. Nature Genetics 48: 694–699

Daron J, Slotkin RK (2017) EpiTEome: Simultaneous detection of transposable element insertion sites and their DNA methylation levels. Genome Biol 18: 91

De Santa F, Totaro MG, Prosperini E, Notarbartolo S, Testa G, Natoli G (2007) The Histone H3 Lysine-27 Demethylase Jmjd3 Links Inflammation to Inhibition of Polycomb-Mediated Gene Silencing. Cell 130: 1083–1094

Deleris A, Stroud H, Bernatavichute Y, Johnson E, Klein G, Schubert D, Jacobsen SE (2012) Loss of the DNA methyltransferase MET1 Induces H3K9 hypermethylation at PcG target genes and redistribution of H3K27 trimethylation to transposons in Arabidopsis thaliana. PLoS Genet 8: e1003062

Derkacheva M, Steinbach Y, Wildhaber T, Mozgova I, Mahrez W, Nanni P, Bischof S, Gruissem W, Hennig L (2013) Arabidopsis MSI1 connects LHP1 to PRC2 complexes. EMBO J 32: 2073–85

Dubin MJ, Mittelsten Scheid O, Becker C (2018) Transposons: a blessing curse. Current Opinion in Plant Biology 42: 23–29

Durr J, Papareddy R, Nakajima K, Gutierrez-Marcos J (2018) Highly efficient heritable targeted deletions of gene clusters and non-coding regulatory regions in Arabidopsis using CRISPR/Cas9. Sci Rep 8: 4443

Ferrari KJ, Scelfo A, Jammula S, Cuomo A, Barozzi I, Stutzer A, Fischle W, Bonaldi T, Pasini D (2014) Polycomb-dependent H3K27me1 and H3K27me2 regulate active transcription and enhancer fidelity. Mol Cell 53: 49–62

Fuchs J, Jovtchev G, Schubert I (2008) The chromosomal distribution of histone methylation marks in gymnosperms differs from that of angiosperms. Chromosome Research 16: 891–898

Gan ES, Xu Y, Wong JY, Goh JG, Sun B, Wee WY, Huang J, Ito T (2014) Jumonji demethylases moderate precocious flowering at elevated temperature via regulation of FLC in Arabidopsis. Nat Commun 5: 5098

Gendrel AV, Lippman Z, Martienssen R, Colot V (2005) Profiling histone modification patterns in plants using genomic tiling microarrays. Nat Methods 2: 213–8

Gu Z, Eils R, Schlesner M, Ishaque N (2018) EnrichedHeatmap: An R/Bioconductor package for comprehensive visualization of genomic signal associations. BMC Genomics 19: 234–234

Hanna CW, Perez-Palacios R, Gahurova L, Schubert M, Krueger F, Biggins L, Andrews S, Colome-Tatche M, Bourc’his D, Dean W, Kelsey G (2019) Endogenous retroviral insertions drive non-canonical imprinting in extra-embryonic tissues. Genome Biol 20: 225

Hou X, Zhou J, Liu C, Liu L, Shen L, Yu H (2014) Nuclear factor Y-mediated H3K27me3 demethylation of the SOC1 locus orchestrates flowering responses of Arabidopsis. Nature Communications 5: 1–14

Ibarra CA, Feng X, Schoft VK, Hsieh TF, Uzawa R, Rodrigues JA, Zemach A, Chumak N, Machlicova A, Nishimura T, Rojas D, Fischer RL, Tamaru H, Zilberman D (2012) Active DNA demethylation in plant companion cells reinforces transposon methylation in gametes. Science 337: 1360–1364

Jacob Y, Feng S, LeBlanc CA, Bernatavichute YV, Stroud H, Cokus S, Johnson LM, Pellegrini M, Jacobsen SE, Michaels SD (2009) ATXR5 and ATXR6 are H3K27 monomethyltransferases required for chromatin structure and gene silencing. Nature Structural & Molecular Biology 16: 763–768

Jacob Y, Stroud H, LeBlanc C, Feng S, Zhuo L, Caro E, Hassel C, Gutierrez C, Michaels SD, Jacobsen SE (2010) Regulation of heterochromatic DNA replication by histone H3 lysine 27 methyltransferases. Nature 466: 987–991

Johannes F, Porcher E, Teixeira FK, Saliba-Colombani V, Simon M, Agier N, Bulski A, Albuisson J, Heredia F, Audigier P, Bouchez D, Dillmann C, Guerche P, Hospital F, Colot V (2009) Assessing the Impact of Transgenerational Epigenetic Variation on Complex Traits. PLoS Genetics 5: e1000530–e1000530

Kakutani T, Jeddeloh JA, Flowers SK, Munakata K, Richards EJ (1996) Developmental abnormalities and epimutations associated with DNA hypomethylation mutations. Proc Natl Acad Sci U S A 93: 12406–11

Kassis JA, Kennison JA, Tamkun JW (2017) Polycomb and trithorax group genes in drosophila. Genetics 206: 1699–1725

Kato M, Takashima K, Kakutani T (2004) Epigenetic Control of CACTA Transposon Mobility in Arabidopsis thaliana. Genetics 168: 961–969

Kleinboelting N, Huep G, Kloetgen A, Viehoever P, Weisshaar B (2012) GABI-Kat SimpleSearch: new features of the Arabidopsis thaliana T-DNA mutant database. Nucleic Acids Res 40: D1211–5

Kolde R (2015) pheatmap: Pretty heatmaps [Software]. In

Kouzarides T (2007) Chromatin Modifications and Their Function. Cell 128: 693–705

Krueger F, Andrews SR (2011) Bismark: a flexible aligner and methylation caller for Bisulfite-Seq applications. Bioinformatics 27: 1571–2

Lafos M, Kroll P, Hohenstatt ML, Thorpe FL, Clarenz O, Schubert D (2011) Dynamic Regulation of H3K27 Trimethylation during Arabidopsis Differentiation. PLoS Genetics 7: e1002040–e1002040

Lan F, Bayliss PE, Rinn JL, Whetstine JR, Wang JK, Chen S, Iwase S, Alpatov R, Issaeva I, Canaani E, Roberts TM, Chang HY, Shi Y (2007) A histone H3 lysine 27 demethylase regulates animal posterior development. Nature 449: 689–694

Langmead B (2010) Aligning short sequencing reads with Bowtie. Curr Protoc Bioinformatics Chapter 11: Unit 11 7

Larsson J (2019) *eulerr*: Area-proportional *Euler* and *Venn* diagrams with ellipses. In

Laugesen A, Hojfeldt JW, Helin K (2019) Molecular Mechanisms Directing PRC2 Recruitment and H3K27 Methylation. Mol Cell 74: 8–18

Lee MG, Villa R, Trojer P, Norman J, Yan KP, Reinberg D, Croce LD, Shiekhattar R (2007) Demethylation of H3K27 Regulates Polycomb Recruitment and H2A Ubiquitination. Science 318: 447–450

Lewis EB (1978) A gene complex controlling segmentation in Drosophila. Nature 276: 565–70

Li C, Gu L, Gao L, Chen Chen C-QWQQC-WCSWLJL-FAC-YCSYVNYQMPSAL, Cui Y (2016) Concerted genomic targeting of H3K27 demethylase REF6 and chromatin-remodeling ATPase BRM in Arabidopsis. Nature genetics 48: 687–693

Liu C, Lu F, Cui X, Cao X (2010) Histone methylation in higher plants. Annu Rev Plant Biol 61: 395–420

Liu J, Feng L, Gu X, Deng X, Qiu Q, Li Q, Zhang Y, Wang M, Deng Y, Wang E, He Y, Bäurle I, Li J, Cao X, He Z (2019) An H3K27me3 demethylase-HSFA2 regulatory loop orchestrates transgenerational thermomemory in Arabidopsis. Cell Research 29: 379–390

Lu F, Cui X, Zhang S, Jenuwein T, Cao X (2011) Arabidopsis REF6 is a histone H3 lysine 27 demethylase. Nature Genetics 43: 715–719

Marí-Ordóñez A, Marchais A, Etcheverry M, Martin A, Colot V, Voinnet O (2013) Reconstructing de novo silencing of an active plant retrotransposon. Nature Genetics 45: 1029–1039

Mathieu O, Reinders J, Caikovski M, Smathajitt C, Paszkowski J (2007) Transgenerational stability of the Arabidopsis epigenome is coordinated by CG methylation. Cell 130: 851–62

Matzke MA, Mosher RA (2014) RNA-directed DNA methylation: an epigenetic pathway of increasing complexity. Nat Rev Genet 15: 394–408

Mirouze M, Lieberman-Lazarovich M, Aversano R, Bucher E, Nicolet J, Reinders J, Paszkowski J (2012) Loss of DNA methylation affects the recombination landscape in Arabidopsis. Proceedings of the National Academy of Sciences 109: 5880–5885

Miura A, Yonebayashi S, Watanabe K, Toyama T, Shimada H, Kakutani T (2001) Mobilization of transposons by a mutation abolishing full DNA methylation in Arabidopsis. Nature 411: 212–4

Molitor A, Latrasse D, Zytnicki M, Andrey P, Houba-Hérin N, Hachet M, Battail C, Del Prete S, Alberti A, Quesneville H, Gaudin V (2016) The Arabidopsis hnRNP-Q Protein LIF2 and the PRC1 subunit LHP1 function in concert to regulate the transcription of stress-responsive genes. The Plant Cell 28: tpc.00244.2016-tpc.00244.2016

Montgomery ND, Yee D, Chen A, Kalantry S, Chamberlain SJ, Otte AP, Magnuson T (2005) The Murine Polycomb Group Protein Eed Is Required for Global Histone H3 Lysine-27 Methylation. Current Biology 15: 942–947

Noh B, Lee S-H, Kim H-J, Yi G, Shin E-A, Lee M, Jung K-J, Doyle MR, Amasino RM, Noh Y-S (2004) Divergent Roles of a Pair of Homologous Jumonji/Zinc-Finger–Class Transcription Factor Proteins in the Regulation of Arabidopsis Flowering Time. The Plant Cell 16: 2601–2613

Oliver JL, Barturen G, Rueda A, Hackenberg M (2014) MethylExtract: High-Quality methylation maps and SNV calling from whole genome bisulfite sequencing data. F1000Research 2: 217–217

Ong-Abdullah M, Ordway JM, Jiang N, Ooi SE, Kok SY, Sarpan N, Azimi N, Hashim AT, Ishak Z, Rosli SK, Malike FA, Bakar NA, Marjuni M, Abdullah N, Yaakub Z, Amiruddin MD, Nookiah R, Singh R, Low ET, Chan KL et al. (2015) Loss of Karma transposon methylation underlies the mantled somaclonal variant of oil palm. Nature 525: 533–7

Pedersen BS, Eyring K, De S, Yang IV, Schwartz DA (2014) Fast and accurate alignment of long bisulfite-seq reads.

Pfluger J, Wagner D (2007) Histone modifications and dynamic regulation of genome accessibility in plants. Current Opinion in Plant Biology 10: 645–652

Qiu Q, Mei H, Deng X, He K, Wu B, Yao Q, Zhang J, Lu F, Ma J, Cao X (2019) DNA methylation repels targeting of Arabidopsis REF6. Nature Communications 10: 2063–2063

Reddington JP, Sproul D, Meehan RR (2014) DNA methylation reprogramming in cancer: does it act by re-configuring the binding landscape of Polycomb repressive complexes? Bioessays 36: 134–40

Roudier F, Ahmed I, Bérard C, Sarazin A, Mary-Huard T, Cortijo S, Bouyer D, Caillieux E, Duvernois-Berthet E, Al-Shikhley L, Giraut L, Desprás B, Drevensek S, Barneche F, Dérozier S, Brunaud V, Aubourg S, Schnittger A, Bowler C, Martin-Magniette ML et al. (2011) Integrative epigenomic mapping defines four main chromatin states in Arabidopsis. EMBO Journal 30: 1928–1938

Saksouk N, Barth TK, Ziegler-Birling C, Olova N, Nowak A, Rey E, Mateos-Langerak J, Urbach S, Reik W, Torres-Padilla ME, Imhof A, Dejardin J, Simboeck E (2014) Redundant mechanisms to form silent chromatin at pericentromeric regions rely on BEND3 and DNA methylation. Mol Cell 56: 580–94

Shen L, Shao NY, Liu X, Maze I, Feng J, Nestler EJ (2013) diffReps: detecting differential chromatin modification sites from ChIP-seq data with biological replicates. PLoS One 8: e65598

Slotkin RK, Vaughn M, Borges F, Tanurdžić M, Becker JD, Feijó JA, Martienssen RA (2009) Epigenetic Reprogramming and Small RNA Silencing of Transposable Elements in Pollen. Cell 136: 461–472

Stroud H, Do T, Du J, Zhong X, Feng S, Johnson L, Patel DJ, Jacobsen SE (2014) Non-CG methylation patterns shape the epigenetic landscape in Arabidopsis. Nat Struct Mol Biol 21: 64–72

Swigut T, Wysocka J (2007) H3K27 Demethylases, at Long Last. Cell 131: 29-32

Thorvaldsdóttir H, Robinson JT, Mesirov JP (2013) Integrative Genomics Viewer (IGV): High-performance genomics data visualization and exploration. Briefings in Bioinformatics 14: 178–192

Tian T, Liu Y, Yan H, You Q, Yi X, Du Z, Xu W, Su Z (2017) AgriGO v2.0: A GO analysis toolkit for the agricultural community, 2017 update. Nucleic Acids Research 45: W122–W129

Vakoc CR, Sachdeva MM, Wang H, Blobel GA (2006) Profile of Histone Lysine Methylation across Transcribed Mammalian Chromatin. Molecular and Cellular Biology 26: 9185–9195

Vongs A, Kakutani T, Martienssen RA, Richards EJ (1993) Arabidopsis thaliana DNA methylation mutants. Science 260: 1926–8

Wang X, Gao J, Gao S, Song Y, Yang Z, Kuai B (2019) The H3K27me3 demethylase REF6 promotes leaf senescence through directly activating major senescence regulatory and functional genes in Arabidopsis. PLOS Genetics 15: e1008068–e1008068

Weinhofer I, Hehenberger E, Roszak P, Hennig L, Kohler C (2010) H3K27me3 profiling of the endosperm implies exclusion of polycomb group protein targeting by DNA methylation. PLoS Genet 6

Wibowo A, Becker C, Durr J, Price J, Spaepen S, Hilton S, Putra H, Papareddy R, Saintain Q, Harvey S, Bending GD, Schulze-Lefert P, Weigel D, Gutierrez-Marcos J (2018) Partial maintenance of organ-specific epigenetic marks during plant asexual reproduction leads to heritable phenotypic variation. Proc Natl Acad Sci U S A 115: E9145–E9152

Yan W, Chen D, Smaczniak C, Engelhorn J, Liu H, Yang W, Graf A, Carles CC, Zhou D-X, Kaufmann K (2018) Dynamic and spatial restriction of Polycomb activity by plant histone demethylases. Nature Plants 4: 681–689

Yu X, Li L, Li L, Guo M, Chory J, Yin Y (2008) Modulation of brassinosteroid-regulated gene expression by jumonji domain-containing proteins ELF6 and REF6 in Arabidopsis. Proceedings of the National Academy of Sciences 105: 7618–7623

Zemach A, Kim MY, Hsieh PH, Coleman-Derr D, Eshed-Williams L, Thao K, Harmer SL, Zilberman D (2013) The Arabidopsis nucleosome remodeler DDM1 allows DNA methyltransferases to access H1-containing heterochromatin. Cell 153: 193–205

Zhang X, Germann S, Blus BJ, Khorasanizadeh S, Gaudin V, Jacobsen SE (2007) The Arabidopsis LHP1 protein colocalizes with histone H3 Lys27 trimethylation. Nature structural & molecular biology 14: 869–71

Zhang Y, Liu T, Meyer CA, Eeckhoute J, Johnson DS, Bernstein BE, Nusbaum C, Myers RM, Brown M, Li W, Liu XS (2008) Model-based analysis of ChIP-Seq (MACS). Genome Biol 9: R137

Zheng S, Hu H, Ren H, Yang Z, Qiu Q, Qi W, Liu X, Chen X, Cui X, Li S, Zhou B, Sun D, Cao X, Du J (2019) The Arabidopsis H3K27me3 demethylase JUMONJI 13 is a temperature and photoperiod dependent flowering repressor. Nature Communications 10: 1303–1303

